# A bioinformatics screen reveals Hox and chromatin remodeling factors at the *Drosophila* histone locus

**DOI:** 10.1101/2023.01.06.523008

**Authors:** Lauren J. Hodkinson, Connor Smith, H. Skye Comstra, Eric H. Albanese, Bukola A. Ajani, Kawsar Arsalan, Alvero Perez Daisson, Katherine B. Forrest, Elijah H. Fox, Matthew R. Guerette, Samia Khan, Madeleine P. Koenig, Shivani Lam, Ava S. Lewandowski, Lauren J. Mahoney, Nasserallah Manai, JonCarlo Miglay, Blake A. Miller, Olivia Milloway, Vu D. Ngo, Nicole F. Oey, Tanya A. Punjani, HaoMin SiMa, Hollis Zeng, Casey A. Schmidt, Leila E. Rieder

**Author notes:** authors contributed equally (Lauren J. Hodkinson, Connor Smith, Casey A. Schmidt, Leila E. Rieder).

## Abstract

Cells orchestrate histone biogenesis with strict temporal and quantitative control. To efficiently regulate histone biogenesis, the repetitive *Drosophila melanogaster* replication-dependent histone genes are arrayed and clustered at a single locus. Regulatory factors concentrate in a nuclear body known as the histone locus body (HLB), which forms around the locus. Historically, HLB factors are largely discovered by chance, and few are known to interact directly with DNA. It is therefore unclear how the histone genes are specifically targeted for unique and coordinated regulation. To expand the list of known HLB factors, we performed a candidate-based screen by mapping 30 publicly available ChIP datasets and 27 factors to the *Drosophila* histone gene array. We identified novel transcription factor candidates, including the *Drosophila* Hox proteins Ultrabithorax, Abdominal-A and Abdominal-B, suggesting a new pathway for these factors in influencing body plan morphogenesis. Additionally, we identified six other transcription factors that target the histone gene array: JIL-1, Hr78, the long isoform of fs(1)h as well as the generalized transcription factors TAF-1, TFIIB, and TFIIF. Our foundational screen provides several candidates for future studies into factors that may influence histone biogenesis. Further, our study emphasizes the powerful reservoir of publicly available datasets, which can be mined as a primary screening technique.

## Introduction

Cells rely on strict temporal and quantitative orchestration of gene expression. One way the nucleus accomplishes coordinated gene regulation is through the establishment of nuclear bodies (NBs), membraneless concentrations of proteins and RNAs. The NB micro-environment facilitates processes such as efficient gene expression through transcription and RNA-processing (Matera *et al*. 2009; Tatomer *et al*. 2016; Arias Escayola and Neugebauer 2018).

The histone locus body (HLB) is a conserved NB that regulates histone gene expression and forms at the loci of the replication-dependent histone genes (Duronio and Marzluff 2017) in many different organisms, including humans and *Drosophila*. The HLB is characterized by a set of factors that collectively regulate the uniquely organized histone genes. The *Drosophila melanogaster* histone locus is a cluster of ~100 tandemly repeated arrays, in which each 5 Kb array includes the 5 canonical histone genes along with their respective promoters and regulatory elements (McKay *et al*. 2015; Duronio and Marzluff 2017; Bongartz and Schloissnig 2018). Each array contains two TATA-box containing promoters, one for *H3* and *H4* and one *H2A* and *H2B*. Additionally, the *H1* gene has its own unique promoter that lacks a TATA-box. These promoters contain some known motifs (Crayton *et al*. 2004; Isogai *et al*. 2007; Rieder *et al*. 2017) that interact with DNA-binding factors to initiate and regulate histone transcription. The clustered, repetitive organization of the locus allows for precise HLB formation at a single genomic location and highly coordinated histone biogenesis linked to S-phase of the cell cycle (Marzluff *et al*. 2002; White *et al*. 2011).

The *Drosophila* HLB is a well-characterized NB that includes several known components that play a role in both the cell cycle regulation of histone gene transcription and the unique processing of histone RNA transcripts. Several proteins involved in the initiation and regulation of histone gene transcription including Chromatin Linked Adaptor for MSL proteins (CLAMP; (Rieder *et al*. 2017)), Multi Sex combs (Mxc (White *et al*. 2011; Yang *et al*. 2014); the *Drosophila* ortholog of human nuclear Nuclear Protein mapped to the Ataxia-Telangiectasia locus (NPAT; (Terzo *et al*. 2015)), FLICE-associated huge protein (FLASH; (Tatomer *et al*. 2016) and Muscle wasted (Mute; (Bulchand *et al*. 2010). Histone mRNA processing is distinct from that of other mRNAs because histone pre-mRNAs lack polyA tails and introns (Duronio and Marzluff 2017). Several known factors are involved in histone mRNA processing and target the histone gene locus including, the U7snRNP, Stem Loop Binding Protein (SLBP), and Lsm11 (Duronio and Marzluff 2017).

It is currently unclear how non-DNA binding factors identify and target the histone locus. The presence of histone mRNA is likely to play a role (Shevtsov and Dundr 2011) as are the presence of *cis* elements within the histone gene array (Salzler *et al*. 2013; Rieder *et al*. 2017). One critical interaction involves CLAMP, a DNA-binding factor that targets loci genome-wide, including the histone gene array by recognizing GA-repeat sequences in the *H3/H4* promoter (Rieder *et al*. 2017). Although the presence of CLAMP is critical for the localization of critical HLB-specific factors such as Mxc (Rieder *et al*. 2017), the interaction between CLAMP and GA-repeat is not always necessary for HLB formation (Koreski *et al*. 2020) and CLAMP is not sufficient for HLB formation (Rieder *et al*. 2017). Therefore, it is likely that other DNA-interacting proteins participate in defining the locus for HLB-specific factors. We still lack a comprehensive list of factors associated with histone biogenesis and therefore the mechanisms of histone gene regulation remain incomplete.

Historically, novel HLB factors are often discovered by chance through immunofluorescence, for example: CLAMP (Rieder *et al*. 2017), Myc (Daneshvar *et al*. 2011), Mute (Bulchand *et al*. 2010), and Abnormal oocyte (Berloco *et al*. 2001). To discover novel DNA-binding proteins that target the histone locus, we used a candidate-based bioinformatics screen. We leveraged publicly available *Drosophila* ChIP-seq data sets and knowledge of histone gene regulation to curate and analyze a list of candidate DNA-binding factors. We used a bioinformatics pipeline on Galaxy (Afgan *et al*. 2016; The Galaxy Community 2022) to map candidate ChIP-seq data to a single copy of the histone gene array. The ~107 histone gene arrays are nearly identical in sequence (Bongartz and Schloissnig 2018) and we can collapse -omics data from the entire locus onto a single array (McKay *et al*. 2015; Rieder *et al*. 2017; Koreski *et al*. 2020). Supervised undergraduate students conducted much of the initial screen as part of a course-based undergraduate research experience (CURE; (Schmidt *et al*. 2022), demonstrating the simplicity and versatility of the pipeline design. We discovered several DNA-interacting proteins that pass our initial bioinformatics screen. Our novel candidates that target the histone gene array include development transcription factors such as Hox factors, which may provide a mechanistic link between segment identity and cell division.

Future wet lab studies are required to confirm the presence of these candidates at the histone locus, determine any tissue and temporal specificity, and describe the precise roles of candidates in HLB formation and histone biogenesis. As a whole, our screen establishes mining of existing -omics data as a tool to identify new candidate HLB factors. Although we are limited by the factors, tissues, treatments, and timepoints interrogated by the dataset generators, our pipeline is an inexpensive and rapid tool to screen candidate factors for future wet-lab study

## Methods and Materials

### GEO Datasets

All data sets were downloaded from the NCBI SRA run selector through the gene expression omnibus (GEO). See Table 1 for Accession numbers and references.

**Table 1:** DNA-binding factor candidate datasets

| Candidate | GEO<br>Accession # | SRA Run Selector # | Paper citation |
| --- | --- | --- | --- |
| **Abd-A**<br>Abdominal-A | GSE69796 | **anti-GFP ChIP DNA from Kc167 cells expressing AbdA-GFP**<br>1 - SRR2060648 2 - SRR2060649<br>**Input**<br>1 - SRR2060652 2 - SRR2060653 | (Beh *et al.* 2016) |
| **Abd-B**<br>Abdominal-B | GSE69796 | **anti-GFP ChIP DNA from Kc167 cells expressing AbdB-GFP**<br>1 - SRR2060650 2 - SRR2060651<br>**Input**<br>1 - SRR2060652 2 - SRR2060653 | (Beh *et al.* 2016) |
| **ANTP**<br>Antennapedia | GSE125604 | **anti-GFP (Invitrogen) from ANTP-GFP genotype**<br>1 - SRR8483063<br>**Input**<br>1 - SRR8483064 | (Kribelbauer *et al.* 2020) |
| **CP190**<br>Centrosomal protein<br>190kD | GSE118699 | **CP190 rabbit (Pai et al 2004)**<br>1 - SRR7706256 2 - SRR7706258<br>**Input**<br>1 - SRR7706251 2 - SRR7706252 | (Bag *et al.* 2019) |
| **CTCF** | GSE175402 | CTCF<br>1 - SRR14631231 2 - SRR14631232<br>**Input**<br>1 - SRR14631233 2 - SRR14631234 | (Kyrchanova *et al.* 2021) |
| **Exd**<br>Extradenticle | GSE125604 | **anti-V5 (Invitrogen) on exd-V5 transgene genotype**<br>1 - SRR8483055<br>**Input**<br>1 - SRR8483056 | (Kribelbauer *et al.* 2020) |
| **Fs(1)h**<br>Female sterile (1)<br>homeotic | GSE42086 | **Female late embryo-derived cell line, ChIP of Fs(1)h long isoform**<br>1 - SRR611533<br>**Female late embryo-derived cell line, ChIP of both isoforms of Fs(1)h**<br>1 - SRR611535<br>**Input**<br>1 - SRR611537 | (Kellner *et al.* 2013) |
| **Gcn5** | GSE83408 | **Gcn5 rabbit polyclonal antibody (5 ug/IP)**<br>1 - SRR3671294 2 - SRR3671295 3 - SRR3671298<br>**Input**<br>1 - SRR3671296 2 - SRR3671297 3 - SRR3671299 | (Ali *et al.* 2017) |
| **Hr78**<br>Hormone-receptor-like 78 | GSE50370 | **Hr78-GFP_8-16_embryonic_ChIP-seq_ChIP**<br>1 - SRR1198798 2 - SRR1198799<br>**Input**<br>1 - SRR1198796 2 - SRR1198797 | (THE MODENCODE<br>CONSORTIUM *et al.* 2010) |
| **Hnf4**<br>Hepatocyte nuclear factor<br>4 | GSE73675 | **rat anti-dHNF4 3600**<br>1 - SRR2548371 2 - SRR2548372<br>3 - SRR2548373 4 - SRR2548374<br>**Inputs**<br>1 - SRR2548367 2 - SRR2548368<br>3 - SRR2548369 4 - SRR2548370 | (Barry and Thummel 2016) |
| **HTH**<br>Homothorax | GSE125604 | **anti-Hth (gp52, N-terminal)**<br>1 - SRR8483065<br>**Input**<br>1 - SRR8483066 | (Kribelbauer *et al.* 2020) |
| **JIL-1** | GSE54438 | **JIL-1 monoclonal antibody 5C9**<br>1 - SRR1145605 2 - SRR1145606<br>**Input**<br>1 - SRR1145612 2 - SRR1145613 | (Cai *et al.* 2014) |
| **M1BP**<br>Motif 1 Binding Protein | GSE97841 | **M1BP_Antibody**<br>1 - SRR10759878<br>**Input**<br>1 - SRR10759877 | (Baumann and Gilmour 2017) |
| **MSL-1**<br>Male-specific Lethal 1 | GSE37864 | **polyclonal rabbit MSL1, crude serum**<br>1 - SRR495366 2 - SRR495367<br>**Input**<br>1 - SRR495378 2 - SRR495380 | (Straub *et al.* 2013) |
| **Ndf/CG4747**<br>Nucleosome-destabilizing<br>factor | GSE42025 | **PAP antibody (Sigma P1291)**<br>1 - SRR611192 2 - SRR611194<br>3 - SRR611196 4 - SRR611198<br>**Input**<br>1 - SRR611193 2 - SRR611195<br>3 - SRR611197 4 - SRR611199 | (Wang *et al.* 2013) |
| **Nej (S2 cells)**<br>Nejire | GSE72666 | **anti-CBP, custom-made antibodies**<br>1 - SRR2232434<br>**Input**<br>1 - SRR2232432 | (Doiguchi *et al.* 2016) |
| **Nej (Embryos)**<br>Nejire | GSE68983 | Nej<br>1 - SRR4044401 | (Koenecke *et al.* 2016) |
|  |  | **Input**<br>1 - SRR2031906 |  |
| **Opa**<br>Odd Paired | GSE140722 | **In-house anti-Opa antibody**<br>1 - SRR10502454 2 - SRR10502455<br>3 - SRR10502458 4 - SRR10502459<br>**Input**<br>1 - SRR10502456 2 - SRR10502457<br>3 - SRR10502460 4 - SRR10502461 | (Koromila *et al.* 2020) |
| **Pan**<br>Pangolin | GSE50340 | **Pan**<br>1 - SRR1198824 2 - SRR1198825<br>**Input**<br>1 - SRR1198822 2 - SRR1198823 | (THE MODENCODE<br>CONSORTIUM *et al.* 2010) |
| **Pnt**<br>Pointed | GSE114092 | **Pnt**<br>1 - SRR7126165<br>**Input**<br>1 - SRR7126164 | (Webber *et al.* 2018) |
| **Psc**<br>Posterior sex combs | GSE38166 | **Psc Mitotic S2**<br>1 - SRR 500149 2 - SRR 500150<br>**Psc Control S2**<br>1 - SRR500151 2 - SRR500152<br>**Psc Mitotic S2 Input**<br>1 - SRR 500153 2 - SRR 500154<br>**Psc Control S2 Input**<br>1 - SRR 500155 2 - SRR 500156 | (Follmer *et al.* 2012) |
| **Scm**<br>Sex comb on midleg | GSE66183 | **BioTAP-N-Scm**<br>1 - SRR1813233 2 - SRR1813243 3 - SRR1813245<br>**Input**<br>1 - SRR1813234 2 - SRR1813244 3 - SRR1813246 | (Kang *et al.* 2015) |
| **su(z)12**<br>suppressor of zeste 12 | GSE36039 | **Su(z)12 ChIP**<br>1 - SRR363407 2 - SRR363408<br>**Input**<br>1 - SRR363409 2 - SRR363410 | (Herz *et al.* 2012) |
| **TAF1**<br>TBP-Associated Factor 1 | GSE97841 | **TAF1 Antibody**<br>1 - SRR5452843 2 - SRR5452844<br>**Inputs**<br>1 - SRR5452847 2 - SRR5452848 | (Baumann and Gilmour 2017) |
| **TFIIB**<br>Transcription Factor II B | GSE120152 | **anti-TFIIB rabbit polyclonal, custom**<br>1 - SRR7874066 2 - SRR7874067<br>**Inputs**<br>1 - SRR7874069 2 - SRR7874070 | (Ramalingam *et al.* 2021) |
| **TFIIF**<br>Transcription Factor II F | GSE120152 | **anti-TFIIF rabbit polyclonal, custom**<br>1 - SRR7874068<br>**Inputs**<br>1 - SRR7874069 | (Ramalingam *et al.* 2021) |
| **TRF2**<br>TBP protein-related factor 2 | GSE97841 | **TRF2 Antibody**<br>1 - SRR5452845 2 - SRR5452846<br>**Inputs**<br>1 - SRR5452847 2 - SRR5452848 | (Baumann and Gilmour 2017) |
| **Ubx (Kc cells)**<br>Ultrabithorax | GSE69796 | **anti-GFP ChIP DNA from Kc167 cells expressing Ubx-GFP**<br>1 - SRR2060646 2 - SRR2060647<br>**Inputs:**<br>1 - SRR2060652 2 - SRR2060653 | (Beh *et al.* 2016) |
| **Ubx (embryos)**<br>Ultrabithorax | GSE64284 | **Anti-V5 ChIP, Ubx-V5**<br>1 - SRR1721317 2 - SRR1721321<br>**Inputs**<br>1 - SRR1721316 2 - SRR1721320 | (Shlyueva *et al.* 2016) |
| **Ubx (larva)**<br>Ultrabithorax | GSE184454 | **Anti-FLAG monoclonal, 3xFLAG-Ubx**<br>1 - SRR15972582 2 - SRR15972584<br>**Inputs**<br>1 - SRR15972583 2 - SRR15972585 | (Feng *et al.* 2022) |

### Bioinformatic Analysis and Data Visualization

We directly imported individual FASTQ data sets into the web-based platform Galaxy (Afgan *et al*. 2016; The Galaxy Community 2022) through the NCBI SRA run selector by selecting the desired runs and utilizing the computing galaxy download feature. We retrieved the FASTQ files were from the SRA using the “faster download” Galaxy command. Because the ~100 histone gene arrays are extremely similar in sequence (Bongartz and Schloissnig 2018), we can collapse ChIP-seq data onto a single histone array (McKay *et al*. 2015; Bongartz and Schloissnig 2018; Koreski *et al*. 2020). We used a custom “genome” that includes a single *Drosophila melanogaster* histone array similar to that in Mckay *et al*. 2015, which we directly uploaded to Galaxy using the “upload data” feature and normalized using the Galaxy command “normalize fasta” specifying an 80 bp line length for the output FASTA. We aligned ChIP-reads to the normalized histone gene array using Bowtie2 (Langmead and Salzberg 2012) to create BAM files using the user built-in index and “very sensitive end-to-end” parameter settings. We converted the BAM files to bigwig files using the “bamCoverage” Galaxy command in which we set the bin size to 1 bp and set the effective genome size to user specified: 5000 bp (approximate size of l histone array). We also mapped relevant input or IgG datasets, and if available we normalized ChIP datasets to input using the “bamCompare” Galaxy command in which we set the bin size to 1 bp. We visualized the Bigwig files using the Integrative Genome Viewer (IGV) (Robinson *et al*. 2011).

## Results

### Validating the bioinformatics pipeline by mapping TATA-associated factors to the histone gene array

We first sought to validate our bioinformatics pipeline through analysis of known histone locus proteins and associated factors. Isogai *et al*. (2007) used immunofluorescence and cell culture ChIP-qPCR assays to demonstrate that the TATA binding protein (TBP)/TFIID complex selectively binds to the *H3/H4* promoter and the *H2A/H2B* promoter, but TBP-related factor 2 (TRF2) targets the promoter of the TATA-less *H1* promoter. We identified a publicly available a TRF2 ChIP-exo dataset from Baumann *et al*. (2017) for TRF2 and used our pipeline to map the data to the histone gene array. ChIP-exo is similar to ChIP-seq but identifies a more complete set of binding locations for a factor with higher resolution than standard ChIP-seq (Rhee and Pugh 2012). We validated that TRF2 localizes to the H1 promoter (**Figure 1A**). Because we were unable to normalize to an input dataset, we compared the TRF2 alignment to an IgG control. The localization of TRF2 to the TATA-less *H1* promoter is consistent with Isogai *et al*. (2007) and is consistent with where a TBP-related factor (TRF) would be expected to bind as they are known to target TATA-less promoters (Wang *et al*. 2013). Baumann *et al*. 2017 demonstrated that Motif 1 binding protein (M1BP) interacts with TRF2 but that this interaction is mostly restricted to the ribosomal protein (RP) genes (Baumann and Gilmour 2017). We mapped ChIP-exo data for M1BP and observed that it did not localize to the *H1* promoter as we saw with TRF2 nor to any other part of the histone array (**Figure 1A**), further validating our pipeline.

**Figure 1:**
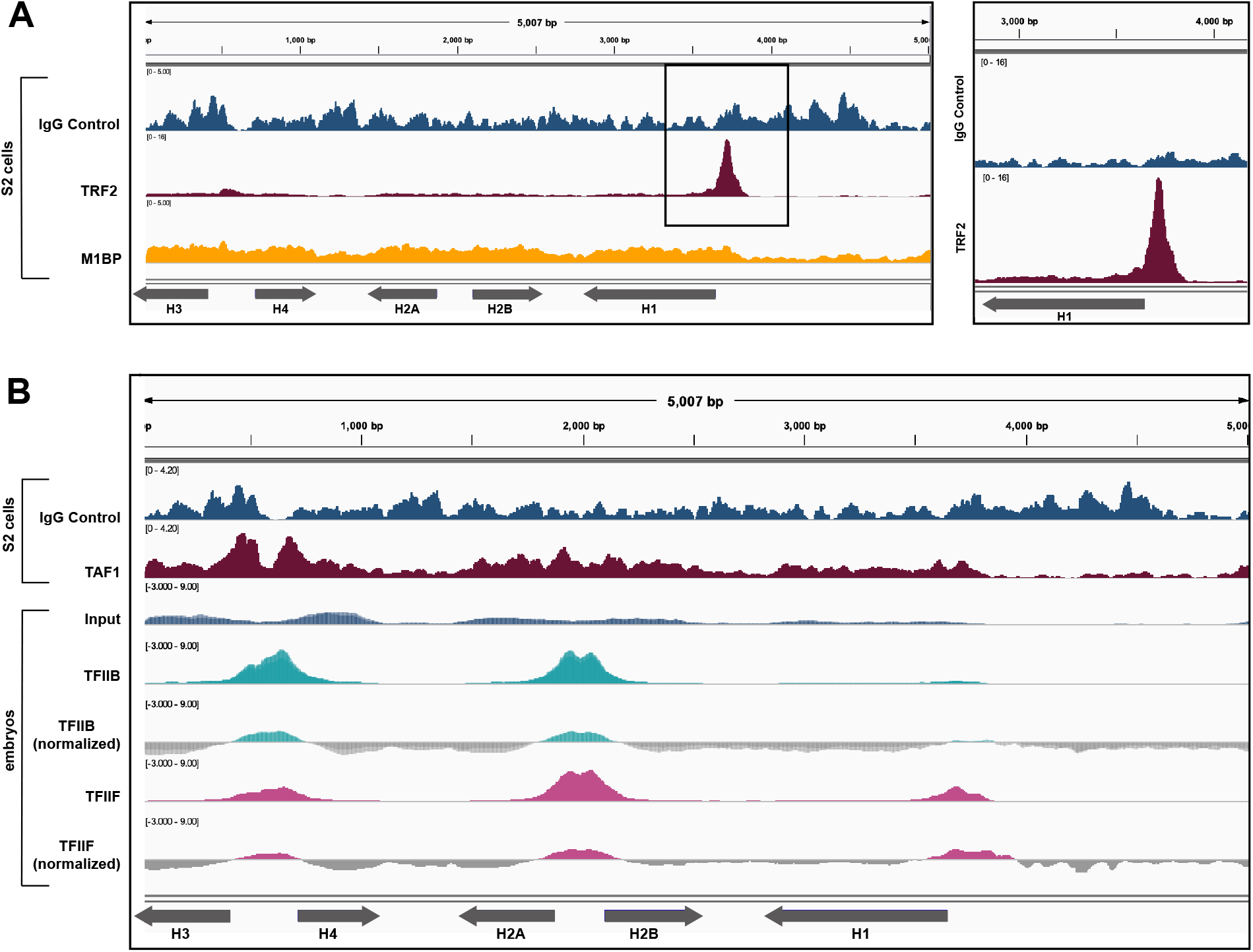
Expected generalized transcription factors localize to the histone array. (A) We mapped ChIP-exo data for TRF2 (maroon, Baumann *et al*. 2017) from S2 cells was aligned to the histone gene array which recapitulates results from Isogia *et al*. 2007 showing localization specifically to the H1 promoter validating our bioinformatics pipeline. We also mapped ChIP-exo data for M1BP (yellow, Baumann *et al*. 2017) which did not localize to the histone gene arry further validating our pipeline ChIP-exo data was compared to an IgG control (blue, Baumann *et al*. 2017 did not provide input sample). (B) We aligned ChIP-exo data for TAF-1 (maroon, Baumann *et al*. 2017) from S2 cells to the histone gene array and compared to a corresponding IgG control. We aligned ChIP-seq datasets for TFIIB (teal, two replicates overlayed, Ramalingam *et al*. 2021) and TFIIF (pink, one replicate, Ramalingam *et al*. 2021) from OregonR mixed populations embryos to the histone gene array and normalized to the provided input signal (blue). TFIIB shows localization to the *H3/H4* promoter and the *H2A/H2B* promoter and TFIIF shows localization to both core promoters and the *H1* promoter confirming that our bioinformatics pipeline can be used to identify novel factors that localize to the histone gene array.

#### Novel general transcription factors that target the histone locus

To expand the list of generalized transcription factors that target the histone locus, we mapped an additional ChIP-exo dataset from Baumann et al. 2017 for TAF1 (TBP associated factor 1). TAF1 is a member of the Transcription Factor IID (TFIID) complex which Isogai *et al* (2007) also suggested localized to the same regions of the histone gene array as TBP. When we mapped the TAF1 ChIP-exo data we observed that TAF1 localizes to the TATA-box regions of the *H3* and *H4* genes and, less robustly, to the TATA-box regions of the *H2A* and *H2B* promoter (**Figure 1B**). Again, we compared this alignment to an IgG control because we were unable to normalize to an input, but because TAF1 associates with TBP which binds to AT rich (TATA box) regions (Baumann and Gilmour 2017), the localization of TAF1 to the TATA-box regions of the core histone genes is expected.

To test the ability of our pipeline to identify novel factors that localize to the histone gene array, we investigated the relationships of additional general transcription factors relationship with the histone array. We identified ChIP-seq datasets for both TFIIB and TFIIF. Both TFIIB and TFIIF are associated with TBP (Ramalingam *et al*. 2021) and therefore we would expect them to localize to the *H3/H4* promoter and *H2A/H2B* promoters, similar to TBP (Isogai *et al*. 2007). We observed both TFIIB and TFIIF localized to the *H3/H4* promoter, *H2A/H2B* promoter while TFIIF also localized, surprisingly, the *H1* promoter (**Figure 1B**).

### Candidate DNA-binding factors that did not pass the bioinformatics screen

After verifying our bioinformatics pipeline, we curated a list of candidate DNA-binding factors (**Table 1** Supplementary Table 1) that we hypothesized would target the histone gene array. To create this candidate list, we prioritized factors that meet at least one of the following criteria: 1) DNA-binding factors with a relationship to a validated HLB factor; 2) DNA-binding factors involved in dosage compensation because CLAMP, a non-sex specific dosage compensation factor, targets the histone locus (Rieder *et al*. 2017; Koreski *et al*. 2020); 3) chromatin remodeling or histone-interacting factors since the epigenetic landscape of the histone locus is largely undefined; 4) early developmental transcription factors since histone gene regulation is critical during early development and synchronized cell division (Chari *et al*. 2019). We also utilized the online platform STRING (Szklarczyk *et al*. 2019) that provides the known and inferred interactomes of a given protein to identify candidates that met the above criteria. Out of the 27 candidates, we rejected 19 as likely not targeting the histone gene array based on the datasets we analyzed.

#### HLB factor-associated candidates

We investigated the DNA-binding factor Sex comb on midleg (Scm), because of its suspected interaction with the known HLB factor Multi-sex combs (Mxc; (White *et al*. 2011; Yang *et al*. 2014). Based on STRING, Scm is predicted to interact with Mxc, as determined by a genetic interference assay in which a double Mxc/Scm mutant resulted in enhanced mutant sex combs phenotypes (Docquier *et al*. 1996; Saget *et al*. 1998). Despite possible interaction with Mxc, neither Scm ChIP-seq data from S2 cells nor 12-24 hr embryos gave meaningful signal over the histone gene array (**Figure 2A**). This result was surprising because the human ortholog of Mxc associates exclusively with the histone promoters (Kaya-Okur *et al*. 2019).

**Figure 2:**
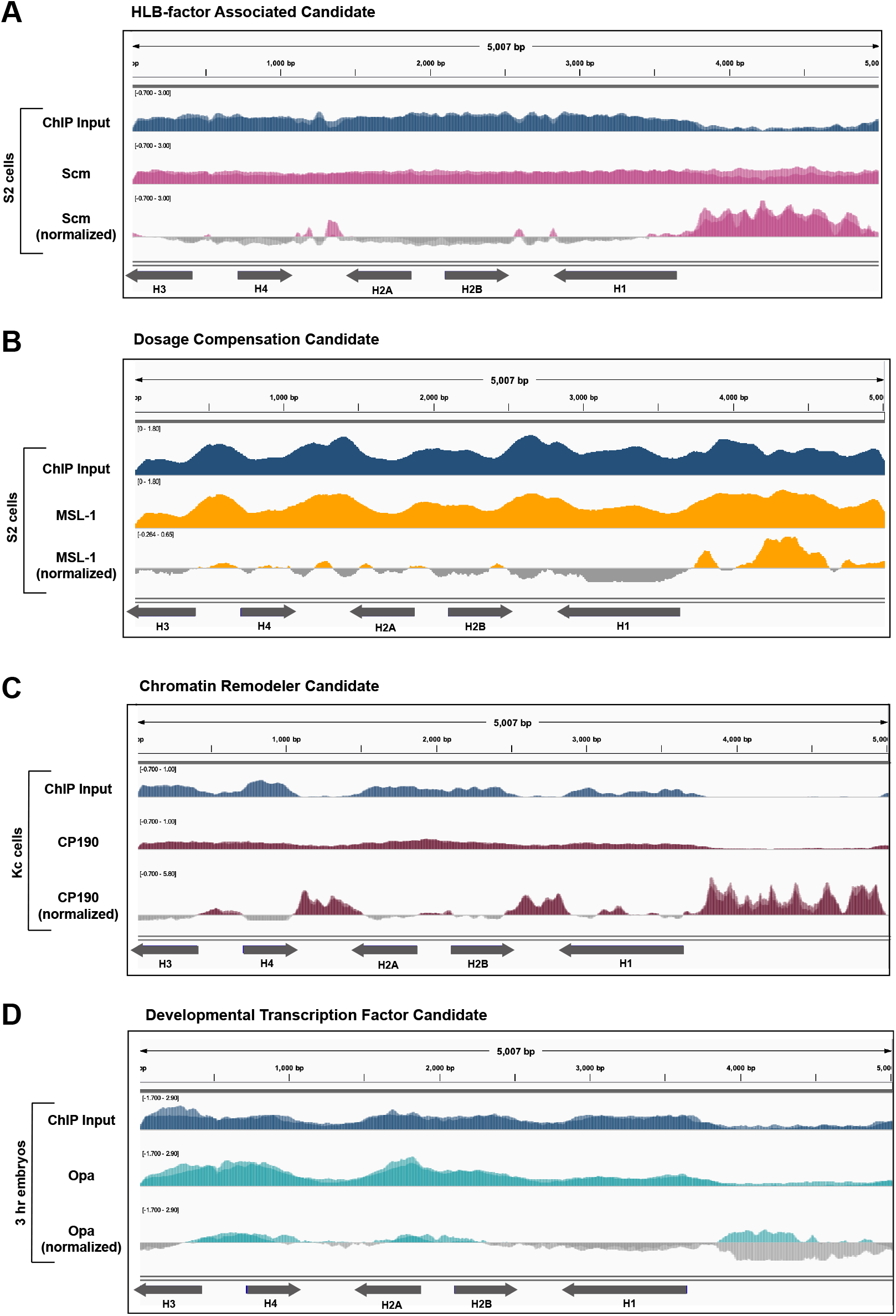
DNA-binding factors from different categories that did not pass the bioinformatics screen. We aligned ChIP-seq datasets for (A) Scm (pink, two replicates overlayed, Kang *et al*. 2015) from S2 cells, (B) MSL-1 (yellow, one replicate, Straub *et al*. 2013) from S2 cells, (C) CP190 (maroon, two replicates overlayed, Bag *et al*. 2019) from Kc cells, and (D) Opa (teal, two replicates overlayed, Koromila *et al*. 2020) from 3 hr mixed population embryos were each aligned to the histone array. Each ChIP signal was normalized to its respective ChIP input signal (blue).

#### Dosage compensation candidates

The HLB factor CLAMP targets the *H3/H4* promoter and regulates histone gene expression (Rieder et al 2017), but also plays additional roles in *Drosophila* male dosage compensation: it binds to GA-rich elements along the male X-chromosome and recruits the Male Specific Lethal complex (MSLc). Further, MSL2, the male specific component of MSLc, also emerged from a cell-based HLB factor screen (White *et al*. 2011) and we recently discovered that MSLc targets one histone gene locus in *Drosophila virilis* (Xie *et al*. 2022b). We therefore hypothesized that dosage compensation factors target the histone gene array along with CLAMP. We chose the following DNA-binding factors for our candidate screen because of their relationship to dosage compensation: MSL1, a protein that scaffolds MSLc (Larschan *et al*. 2006; Straub *et al*. 2013), and nucleosome destabilizing factor (Ndf, CG4747), a putative H3K36me3-binding protein that is important for MSLc localization (Wang *et al*. 2013). When we mapped ChIP-seq datasets from these factors, we found that neither gave meaningful signal over the histone gene array (MSL1 **Figure 2B**, data not shown). This is not surprising as we previously determined that MSL2 does not target the histone locus in *Drosophila melanogaster* by polytene chromosome immunofluorescence (Xie *et al*. 2022b).

#### Chromatin remodeling candidates

One of the lesser-studied characteristics of the histone locus is the regional chromatin environment. The endogenous histone locus is located on chromosome 2L, proximal to pericentric heterochromatin. Despite this proximity, histone expression rapidly increases at the start of G1 in preparation for DNA synthesis during S phase, and quickly ceases upon G2 (Duronio and Marzluff 2017) indicating that chromatin remodeling is likely critical in precisely controlling histone gene expression. We therefore hypothesized that chromatin remodeling factors localize to the histone locus. We chose the following candidates because of their association with chromatin or role in chromatin remodeling: centrosomal 190 kDa protein (CP190), an insulator protein that impacts enhancer protein interactions and stops the spread of heterochromatin (Bag *et al*. 2019); Gcn5, a lysine acetyltransferase critical for oogenesis and morphogenesis (Ali *et al*. 2017); CCCTC-binding factor (CTCF), a genome architectural protein (Kyrchanova *et al*. 2021); Posterior sex combs (Psc), a polycomb-group gene (Follmer *et al*. 2012); and Suppressor 12 of zeste (su(z)12), a subunit of polycomb repressive complex 2 (Herz *et al*. 2012).

After identifying relevant ChIP-seq datasets (**Table 1**), we used our analysis pipeline to map data to the histone gene array. We observed that none of the above chromatin remodeling candidates gave meaningful signal over the histone gene array (CP190 **Figure 2C**, data not shown). We were especially surprised that CP190 did not target the histone array. CP190 binds promoter regions, aids enhancer-promoter interactions, and halts the spreading of heterochromatin. Because the histone locus is proximal to pericentric heterochromatin, we hypothesized the presence of CP190 could explain how centromeric heterochromatin does not expand into the histone locus. In addition, CP190 is a member of the Late Boundary Complex (LBC) (Wolle *et al*. 2015), which also contains the CLAMP protein (Kaye *et al*. 2018). We discovered that the LBC binds to the *H3/H4* promoter region *in vitro* (Xie *et al*. 2022b). We were therefore surprised that CP190 does not appear to target the histone gene array, based on the ChIP-seq datasets we analyzed.

#### Developmental transcription factor candidates

Zygotic histone biogenesis is critical for the constantly dividing embryo; increased histone expression can lengthen the cell cycle while decreased histone biogenesis can shorten the cell cycle (Amodeo *et al*. 2015; Chari *et al*. 2019). Histone biogenesis is tightly coupled to DNA replication, and excess histones are buffered so as not to interfere with zygotic chromatin (Li *et al*. 2012, 2014; Stephenson *et al*. 2021). We therefore hypothesized that early embryonic transcription factors target the histone locus. We chose the following DNA-binding factors based on their roles in the early embryo: Odd pained (Opa), a pair ruled gene that contributes to morphogenesis (Koromila *et al*. 2020); Motif 1 binding protein (M1BP), a transcription pausing factor that interacts with the Hox proteins (Baumann and Gilmour 2017; Bag *et al*. 2021); Hepatocyte nuclear factor 4 (HNF4), a general developmental transcription factor (Barry and Thummel 2016), Pangolin (Pan), a component of the Wingless signaling pathway (Ravindranath and Cadigan 2014); and Pointed (Pnt), a generalized factors the regulates cell proliferation and differentiation in development (Webber *et al*. 2018; Vivekanand 2018). When we mapped appropriate ChIP-seq datasets from these factors, none gave meaningful signal over the histone array (Opa **Figure 2D**, data not shown, see **Figure 1A** for M1BP).

### Candidates that passed the bioinformatics screen

We found that several factors that exhibited distinct, meaningful localization patterns to the histone gene array and therefore warrant further investigation (**Figure 3**). First, we used our bioinformatics pipeline to map a ChIP-seq dataset for the kinase JIL-1, which is responsible for phosphorylating serine 10 on histone 3 (Cai *et al*. 2014; Albig *et al*. 2019). We observed JIL-1 localizing to the histone gene array, specifically to the *H2A/H2B* promoter (**Figure 3A**). We observed an additional sharp peak at the *H3/H4* promoter, but this peak is likely an artifact of short read lengths from the dataset and overlaps with a perfect, long GA-repeat sequence in the *H3/H4* promoter. JIL-1 is a DNA-binding factor that associates with the Maleless helicase (MLE) and MSL-1, two members of MSLc (Albig *et al*. 2019). In addition to CLAMP performing a role in histone biogenesis, it also plays a role in dosage compensation and associates with the MSLc (Larschan *et al*. 2012).

**Figure 3:**
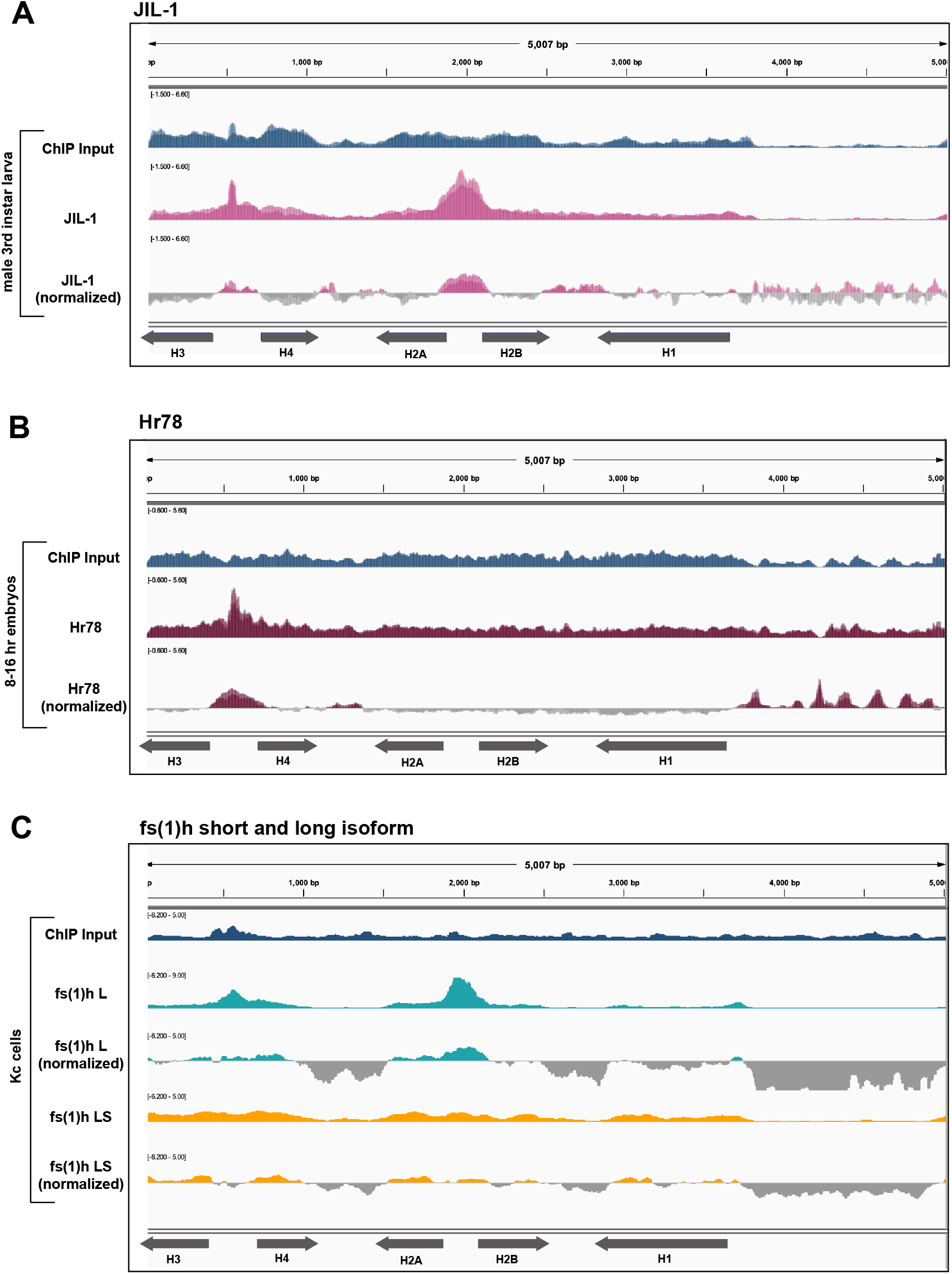
JIL-1, Hr78, and Fs(1)hL localize to the histone gene array. We mapped ChIP datasets for (A) JIL-1 (pink, two replicates overlayed, Cai *et al*. 2014) from male third instar larva, (B) Hr78 (maroon, two replicates overlayed, The MODENCODE Consortium *et al*. 2010) from 8-16 hr mixed population embryos and (C) the long((L, teal) and short (S, yellow) isoform of fs(1)h from Kc cells (Kellner et al. 2013) were all individually aligned to the histone gene array. We normalized each ChIP-seq dataset to its respective input signal (blue).

We also observed hormone-like receptor 78 (Hr78) localize to the *H3-H4* promoter (**Figure 3B**). Finally, we mapped two isoforms of female sterile (1) homeotic (fs(1)h; the *Drosophila* homolog of BRD4). The long and short isoforms of fs(1)h have distinct binding profiles but are assumed have a role in chromatin architecture (Kellner *et al*. 2013). We observed that the long isoform, but not the short isoform, localizes to both the *H2A/H2B* and the *H3/H4* promoters (**Figure 3C**). Interestingly, Kellner *et al*. (013) inferred that the fs(1)h long isoform has a unique role in chromatin remodeling by interacting with specific insulator proteins, one of which is CP190, which did not pass our screen (**Figure 2C**).

### Hox factors localize to the *Drosophila* histone gene array when overexpressed in cell culture

Hox factors are critical for developmental processes like morphogenesis in which cells are constantly dividing and therefore require a near constant supply of histones (Duronio and Marzluff 2017). Histone biogenesis is critical within the first few hours of *Drosophila* development (Amodeo *et al*. 2015; Chari *et al*. 2019). We therefore investigated histone array localization patterns of transcription factors that are critical during early development, including Hox proteins. We identified a publicly available dataset (**Table 1**) in which Beh *et al*. (2016) individually expressed the three Bithorax complex Hox proteins, Ultrabithorax (Ubx), Abdominal-A (Abd-A) and Abdominal-B (Abd-B), in Kc167 cells and performed ChIP-seq. We used our analysis pipeline to map the Ubx, Abd-A, and Abd-B ChIP-seq datasets to the histone gene array and observed striking localization to the *H3/H4* promoter (**Figure 4**). We conclude that when overexpressed in cultured cells, Ubx, Abd-A, and Abd-B all target the histone gene array by ChIP-seq.

**Figure 4:**
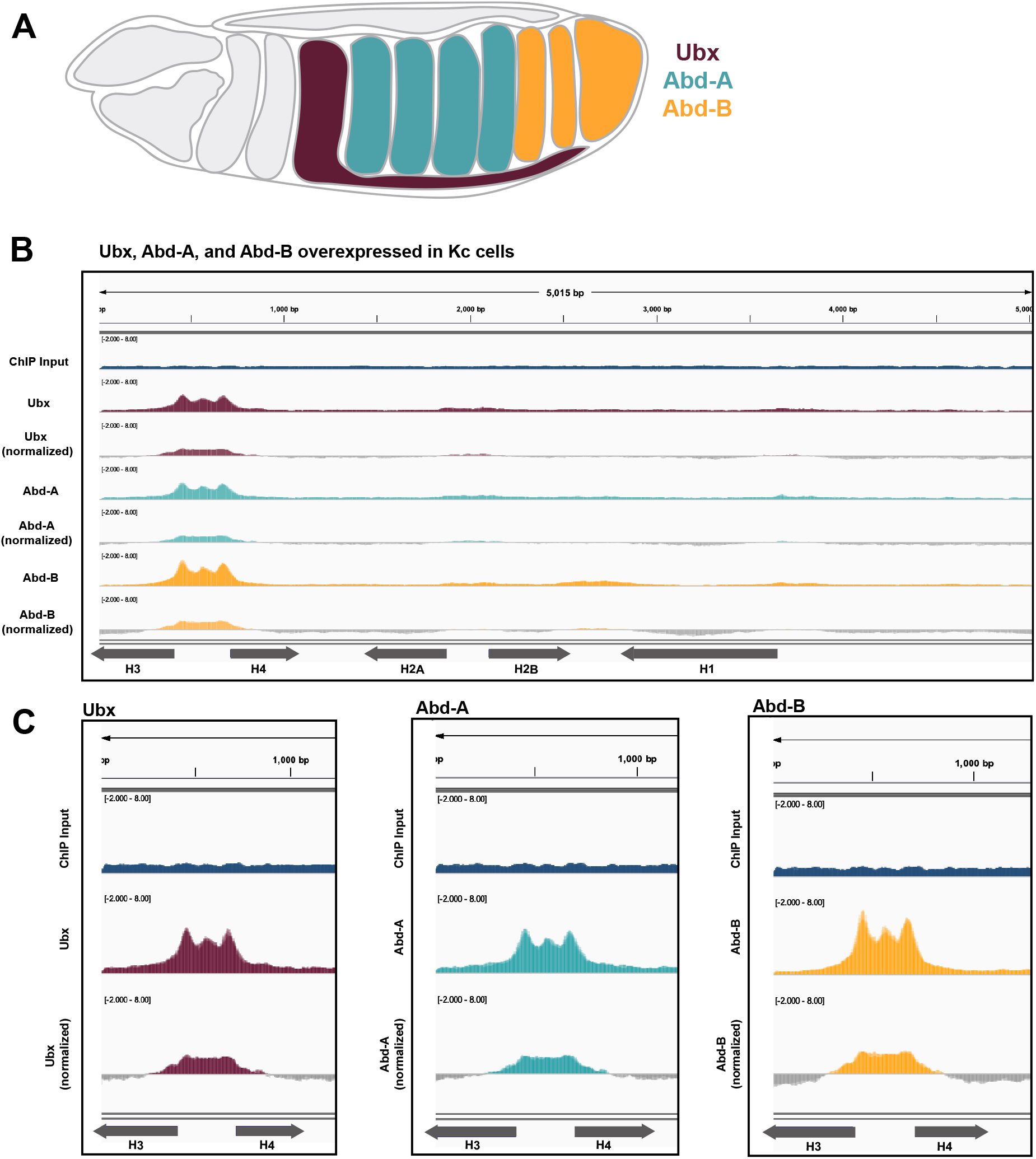
Hox factors Ubx, Abd-A, and Abd-B localize to the histone array. (A) Diagram of relative tissue expression patterns for Ubx (maroon), Abd-A (teal) and Abd-B (yellow). (B) We aligned ChIP-seq datasets from Kc cells expressing Ubx (marron, two replicates overlayed, Beh *et al*. 2016), Abd-A (teal, two replicates overlayed, Beh *et al*. 2016), and Abd-B (yellow, two replicates overlayed, Beh *et al*. 2016) to the histone gene array. We normalized each ChIP-seq dataset to the provided input (blue, two replicates overlayed, Beh *et al*. 2016). (C) Enlarged ChIP-seq signal from (B) of Ubx (maroon), Abd-A (teal), and Abd-B (yellow) over the *H3/H4* promoter.

Because our Hox factor observation (**Figure 4**), could be an artifact of overexpression in cultured cells, we identified two additional Ubx ChIP-seq datasets from 0-16 hr embryos and third instar larval imaginal discs (**Table 1**). We used our pipeline to map these data to the histone gene array and observed that Ubx targets the *H3/H4* promoter and, to a lesser extent, the *H2A/H2B* promoter (**Figure 5**). We conclude that Ubx targets the histone gene array at various developmental stages and in various tissues and is therefore a promising candidate for future wet-lab research designed to validate these bioinformatic observations.

**Figure 5:**
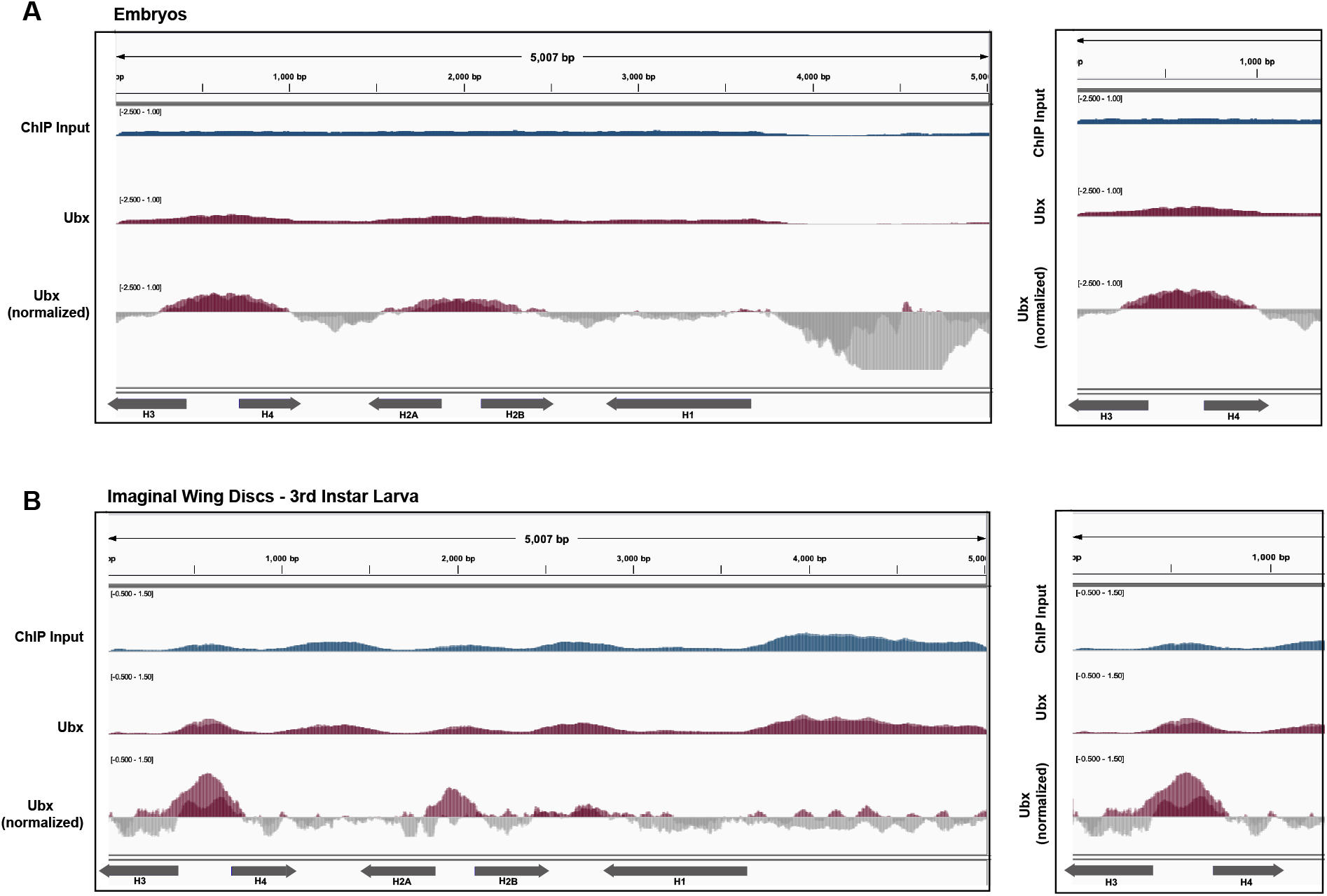
Ubx localizes to the *H3/H4* promoter in embryos and 3^rd^ instar larva. We mapped Ubx ChIP-seq datasets from (A) mixed population embryos (maroon, top panel, two replicates overlayed, Shlyueva *et al*. 2016) and (B) imaginal wing discs in third instar larva (maroon, bottom panel, two replicates overlayed, Feng *et al*. 2022) to the histone gene array. We normalized ChIP-seq datasets to the provided inputs (blue, two replicates overlayed). Signal from the *H3/H4* promoter is enlarged in the panels on the right.

To further investigate the relationship between Hox factors and the histone locus, we identified three additional datasets for Hox proteins and Hox cofactors. There are two different Hox gene complexes in *Drosophila*: the Bithorax complex (which includes Ubx, Abd-A, and Abd-B) and the Antennapedia complex. We first mapped ChIP-seq data for Antennapedia (Antp) (Kribelbauer *et al*. 2020) but did not observe robust localization to the histone gene array (data not shown). We next mapped ChIP-seq data sets for the Hox cofactors extradenticle (Exd) and Homothorax (Hth) (Kribelbauer *et al*. 2020). Exd and Hth associate with the hexapeptide motif in Hox proteins and form heterodimers to impact Hox binding specificity to their gene targets (Rezsohazy *et al*. 2015; Beh *et al*. 2016). We observed that neither Exd nor Hth gave meaningful ChIP signal over the histone gene array (data not shown).

### Power and limitations of the screen

The range of results from our candidate screen demonstrates both the power and limitations of our bioinformatics pipeline. In total, we analyzed datasets for 27 different DNA-binding factors and produced 9 candidates that warrant further wet lab investigation. Despite the power of this screen, we are limited by the availability of public datasets. Characteristics of these datasets, such as quality of reads, read length, and inclusions of controls such as inputs are based on the original experimental design and researchers. Furthermore, we are also restricted by the tissues or genotypes investigated in the original study, limiting the scope of our investigation.

For example, we analyzed several datasets for Nejire (Nej; homolog of mammalian CREB-binding protein (CBP) and Pointed (Pnt). A previous screen in S2 cells identified Nej and Pnt as potential HLB factors (White *et al*. 2011). We mapped a Pnt ChIP-seq dataset from Stage 11 embryos (Table 1) and observed that Pnt does not give meaningful signal over the histone gene array (**Figure 6A**, bottom). Additionally, we investigated two Nej ChIP-seq datasets in which we obtained disparate results. The Nej ChIP-seq dataset from S2 cells did not yield meaningful signal over the histone gene array (**Figure 6A**, center). In contrast, we investigated a Nej ChIP-seq dataset from early *Drosophila* embryos and observed robust localization to the *H3/H4* promoter, *H2A/H2B* promoter and, to a lesser extent, the *H1* promoter (**Figure 6A** top, **Figure 6B**). From these observations, we conclude that Nej likely targets the histone gene array in embryos and would therefore be a strong candidate for future wet-lab studies to validate this observation. Our Pnt and Nej observations demonstrate how our screening approach is powerful but limited by data availability and experiment variables.

**Figure 6:**
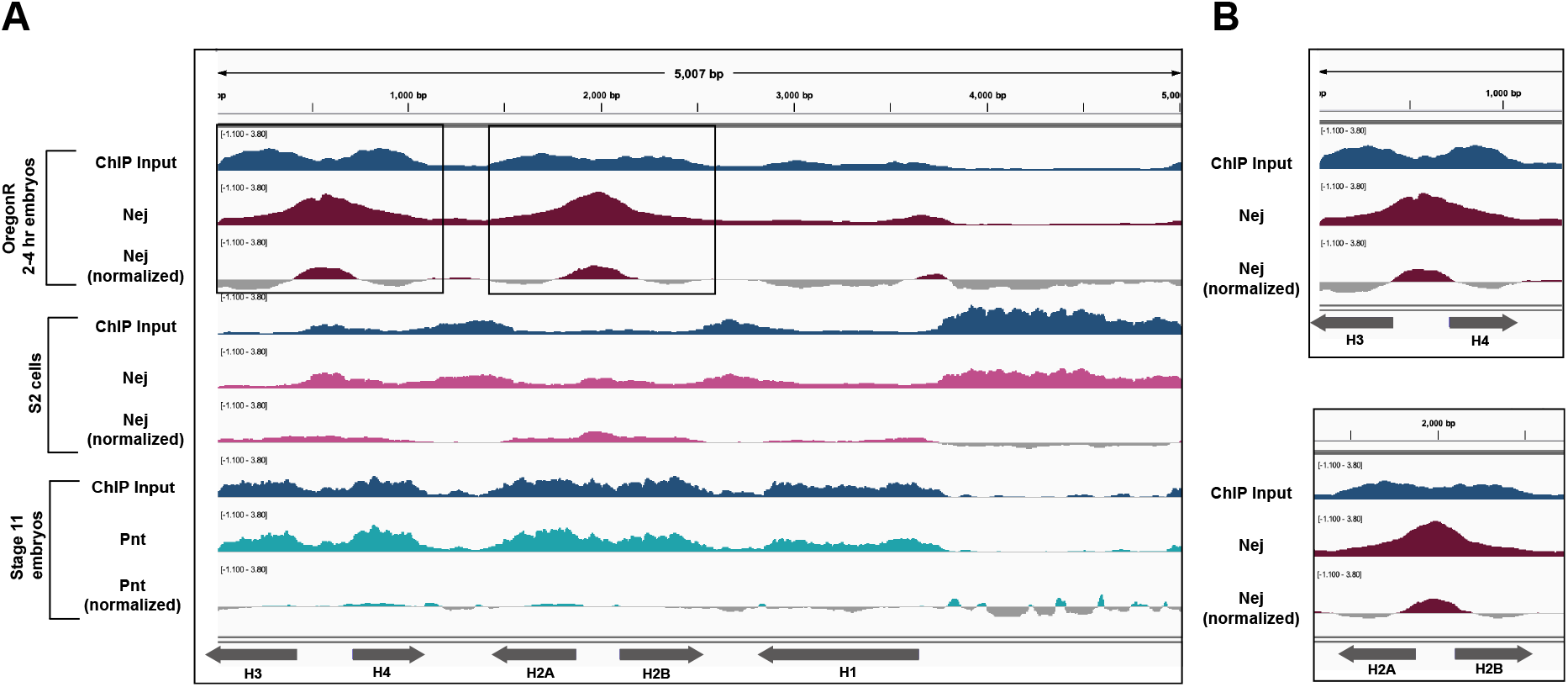
ChIP-seq datasets from different tissues can show different alignment results. We mapped two different ChIP-seq datasets for Nejire (Nej) were aligned to the histone gene array. ChIP data from 2-4 hr embryos (maroon, one replicate, Koenecke et al. 2016), showed localization to the *H/-H4* promoter and the *H2A/H2B* promoter (enlarged in B), while ChIP-seq data from S2 cells (pink, one replicate, Doiguchi *et al*. 2016) showed no localization to the histone gene array. We also aligned ChIP-seq data for Pnt from stage 11 embryos (Webber et al. 2018) to the histone gene array. We normalized the ChIP-seq signals to their respective input signals (blue).

## Discussion

To broaden our understanding of factors that impact histone biogenesis in *Drosophila melanogaster*, we conducted a candidate-based bioinformatics screen for DNA-binding factors that localize to histone gene array. Although many HLB factors are known, it is likely that there are many other factors critical for histone biogenesis that have yet to be identified, since several have been discovered by chance in the past few years including CLAMP (Rieder *et al*. 2017), Winged-Eye (WGE; (Ozawa *et al*. 2016), and Myc (Daneshvar *et al*. 2011). To begin to close this gap in knowledge, we chose 27 factors based on their roles in chromatin remodeling, dosage compensation, development, and interaction with known HLB factors, hypothesizing that these represent strong candidates for novel HLB factors. As our screen is limited by availability of relevant datasets, it will likely produce both false positives and negatives. We therefore envision that the final 9 candidates will be validated through future wet lab experiments (Salzler *et al*. 2013; Rieder *et al*. 2017; Xie *et al*. 2022a).

We validated our bioinformatics pipeline by investigating TRF2, a general transcription factor known to target the histone genes (Isogai *et al*. 2007). We confirmed that TRF2 binds to the TATA-less H1 promoter. Isogai et al. (2007) determined that TBP targets the TATA-containing *H3/H4* and *H2a/H2b* promoters. We expanded this observation by investigating TBP-associated factors TAF1, TFIID, and TFIIF. We discovered that all of these general transcription factors target the histone gene array, further validating our pipeline. We also discovered that the localization of some factors such as Nej to the histone gene array is tissue specific. Nej emerged from a proteomic screen for factors involved in HLB activation in cultured cells (White *et al*. 2011). However, Nej ChIP-seq from cultured cells did not give meaningful signal over the histone gene array, whereas embryo ChIP-seq showed Nej at histone promoters. These observations denote limitations of our screening technique: we are hindered by the availability and quality of datasets for candidate proteins in specific tissues, genotypes, and conditions. Even with the constraints of data availability, we identified 9 out of 27 candidates that give meaningful signal over the histone gene array and warrant future wet lab study.

Many strong candidate factors did not give meaningful ChIP-seq signal over the histone gene array including factors like Scm, which may interact with the confirmed HLB scaffolding factor, Mxc (Docquier *et al*. 1996; Saget *et al*. 1998; Kemp *et al*. 2021). We also investigated factors involved in dosage compensation, including MSL1, Ndf (CG4747), and JIL-1, since the HLB factor CLAMP plays a key role in male X-chromosome activation. MSL2 is a candidate from an unbiased proteomics-based HLB candidate screen in cultured cells (White *et al*. 2011), and we recently discovered that MSLc targets one of the two histone loci in *Drosophila virilis* in salivary gland polytene chromosomes (Xie *et al*. 2022b). Although neither MSL1 nor Ndf localized to the histone gene array, JIL-1 robustly localized to the histone gene array. Of note, the ChIP-seq datasets for MSL1 derived from S2 cells, the Ndf datasets were from both male and female larvae, and the JIL-1 dataset came from specifically male third instar larva. MSL1 and Ndf may target the histone gene array in other tissues or only in embryos, representing potential false negatives. However, JIL-1 is a more generalized kinase that is responsible for phosphorylating serine 10 in histone 3 across the genome, not just on the male X-chromosome (Regnard *et al*. 2011; Cai *et al*. 2014; Albig *et al*. 2019). JIL-1 may therefore be present at the histone locus independent of its role in dosage compensation by contributing to the epigenetic landscape of the histone locus. Taken together, our results indicate that dosage compensation and histone gene expression are likely distinct regulatory events, and the majority of factors are not shared between these processes in *Drosophila melanogaster*.

One of the lesser studied characteristics of the histone locus is the chromatin environment and how epigenetics influences histone gene expression. We identified CP190, Gcn5, Psc, Pangolin, and su(z)12 as chromatin remodeling candidates that might target the histone genes but, after mapping relevant datasets, none of these candidate chromatin remodelers target the histone gene array. We did, however, discover that the long isoform of fs(1)h (fs(1)hL) robustly localizes to the histone gene array. Fs(1)hL has a unique role in chromatin remodeling that differs the short fs(1)h isoform, as it associates with insulator proteins, including CP190 (Kellner *et al*. 2013). Since the histone locus is situated near heterochromatin, it is possible that insulators prevent spreading of heterochromatin into the histone locus. CP190 was also a strong candidate for histone locus-association. CLAMP and CP190 share binding profiles at many promoters and each is important for the other’s localization (Bag *et al*. 2019). However, when we mapped a CP190 ChIP-seq dataset from female embryos, we did not observe histone array localization. Based on these observations, we conclude that fs(1)hL is a strong candidate for future wet lab studies. Fs(1)hL and CLAMP may interact with CP190 at the histone locus, in specific tissues or at precise developmental timepoints.

Finally, we explored several developmental transcription factors because histone biogenesis is critical in the first few hours of *Drosophila* development during rapid zygotic rapid cell divisions. We chose Opa, M1BP, and HNF4 as candidates. Despite their roles in early development, these factors did not target the histone gene array. However, we identified Nej (CREB-binding protein; CBP) as a candidate that targets the histone gene array, specifically in *Drosophila* embryos but not in S2 cells. Nej was previously identified as an HLB candidate through a cell-based proteomics screen (White *et al*. 2011). Nej is a histone acetyltransferase, but it has roles in cell proliferation and developmental patterning. Nej could influence the chromatin environment of the histone locus during key times in development or in tissues that are constantly dividing where histone proteins would be needed. Because of the roles Nej plays in general developmental processes, it is a strong candidate for future wet lab studies.

We were surprised to discover that the Hox proteins Ubx, Abd-A and Abd-B, all localize to the histone array when overexpressed in Kc cells. Specifically, these factors all target the *H3/H4* promoter. This ~300 bp promoter is unique within the 5 Kb histone gene array; it is the minimal sequence required for Mxc localization and HLB formation (Salzler *et al*. 2013) and contains critical GA-repeat *cis* elements targeted by CLAMP (Rieder *et al*. 2017). The CLAMP-GA-repeat interaction promotes recruitment of histone-locus specific transcription factors (Rieder *et al*. 2017; Koreski *et al*. 2020). To confirm that our observations are not a byproduct of Hox overexpression, we also investigated independent Ubx ChIP-seq datasets prepared from early embryos (0-16 hrs) and from third instar larval imaginal wing discs. These datasets confirm that Ubx targets the histone gene array, although the distribution across the array varies between tissues. Ubx, as well as Abd-A and Abd-B, are all highly active in the early embryo when histone proteins are needed to organize newly synthesized DNA. Therefore Ubx, Abd-A, and Abd-B could provide a spatial and temporal link between histone biogenesis, cell division, and morphogenesis in the embryo.

With 9 out of 27 hits from our screen emerging as strong candidates for future studies, our screen has proven to be a powerful tool to identify strong candidates for DNA-binding factors that target this histone gene array. However, our screen also demonstrates the limitations of using publicly available data. Although we curated a list of candidates that were based on known characteristics of histone biogenesis, we were limited by several aspects of these datasets, such as quality of reads, read length, and inclusions of proper controls such as inputs. Controls are specifically important to our pipeline because relative peaks at a given location do not always represent true localization. Our negative data shows a high range of negative signals displayed in **Figure 1**. In some cases, we saw clear enrichment for open chromatin regions, over promoters and/or gene bodies, but did not characterize these as hits. These regions can be overrepresented in the ChIP sequencing experiment as a whole and, therefore, do not reflect where the DNA-binding factor is truly localizing. This is best demonstrated when looking at inputs that also show enrichment over open chromatin or gene bodies as shown in our negative hits figure (**Figure 2**). Inputs between datasets can be highly variable (e.g. **Figure 2, 6**) and, because they are used in the normalization process, can bias the final visualization.

The HLB was discovered by Liu and Gall only fifteen years ago (Liu *et al*. 2006). Since then, novel HLB factors have largely been discovered one at a time by chance. A proteomic screen identified several novel candidates but searched specifically for factors that affect phosphorylation of Mxc in cultured cells (White *et al*. 2011). A comprehensive inventory of HLB factors is necessary to establish a thorough mechanism of histone biogenesis. Histone regulation is especially critical in the early animal embryo: excess histones drive extra, asynchronous mitotic cycles, while depletion of maternal histone deposition accelerates zygotic transcription in *Drosophila* embryos (Chari *et al*. 2019). The timing of important early developmental events such as the mid-blastula transition is influenced by histone to DNA ratios (Amodeo *et al*. 2015). Histone levels also affect pre-mRNA splicing in human cells (Jimeno-González *et al*. 2015), and *H1* isoform loss-of-function mutations are associated with B cell lymphomas (Yusufova *et al*. 2021). Factors that influence histone biogenesis likely contribute to all of these developmental and disease phenotypes.

Here we present a candidate-based screen for novel histone locus-associating factors. Our screen was largely driven by the undergraduate student coauthors in two stages: first we identified strong candidates based on their established or inferred roles, second, we identified and mapped relevant ChIP-seq datasets to the histone gene array. A similar recent bioinformatic screen searched through thousands of datasets and hundreds of hematopoietic transcription factors for those associated with the repetitive mammalian rRNA array. This analysis identified numerous candidate transcription factors but required intensive computational pairwise comparisons and thresholding (Antony *et al*. 2022). We instead chose an informed, narrow list of initial candidates and identified 9 out of 27 for future wet lab studies. Our results not only identify factors that may be involved in histone biogenesis, but also demonstrate the power of a candidate-based bioinformatics screen driven by students.

## Data Availability Statement

The authors affirm that all datasets used in the screen are available on GEO (Gene Expression Omnibus). All GEO accession numbers and runs from the SRA run selector are specified in **Table 1**.

## Acknowledgments

We would like to thank the Emory University students who piloted the project during its earliest stages: Mary Wang, Greg Kimmerer, Mellisa Xie, Dabin Cho, Henrik Torres, Yono Bulis, Edgar Hsieh, Shaariq Khan, Andre Mijacika, Sean Parker, Rohan Ramdeholl, Annalise Weber, and Kelly Yoon. We also thank Nhi Ngo for participation in the project. We thank all the Rieder Lab members for their helpful contributions to project development.

## References

Afgan, E., D. Baker, M. van den Beek, D. Blankenberg, D. Bouvier et al., 2016 The Galaxy platform for accessible, reproducible and collaborative biomedical analyses: 2016 update. Nucleic Acids Res 44: W3–W10.

Albig, C., C. Wang, G. P. Dann, F. Wojcik, T. Schauer et al., 2019 JASPer controls interphase histone H3S10 phosphorylation by chromosomal kinase JIL-1 in Drosophila. Nat Commun 10: 5343.

Ali, T., M. Krüger, S. Bhuju, M. Jarek, M. Bartkuhn et al., 2017 Chromatin binding of Gcn5 in Drosophila is largely mediated by CP190. Nucleic Acids Res 45: 2384–2395.

Amodeo, A. A., D. Jukam, A. F. Straight, and J. M. Skotheim, 2015 Histone titration against the genome sets the DNA-to-cytoplasm threshold for the Xenopus midblastula transition. Proceedings of the National Academy of Sciences 112: E1086–E1095.

Antony, C., S. S. George, J. Blum, P. Somers, C. L. Thorsheim et al., 2022 Control of ribosomal RNA synthesis by hematopoietic transcription factors. Mol Cell 82: 3826–3839.e9.

Arias Escayola, D., and K. M. Neugebauer, 2018 Dynamics and Function of Nuclear Bodies during Embryogenesis. Biochemistry 57: 2462–2469.

Bag, I., S. Chen, L. F. Rosin, Y. Chen, C.-Y. Liu et al., 2021 M1BP cooperates with CP190 to activate transcription at TAD borders and promote chromatin insulator activity. Nat Commun 12: 4170.

Bag, I., R. K. Dale, C. Palmer, and E. P. Lei, 2019 The zinc-finger protein CLAMP promotes gypsy chromatin insulator function in Drosophila. Journal of Cell Science 132: jcs226092.

Barry, W. E., and C. S. Thummel, 2016 The Drosophila HNF4 nuclear receptor promotes glucose-stimulated insulin secretion and mitochondrial function in adults. Elife 5: e11183.

Baumann, D. G., and D. S. Gilmour, 2017 A sequence-specific core promoter-binding transcription factor recruits TRF2 to coordinately transcribe ribosomal protein genes. Nucleic Acids Res 45: 10481–10491.

Beh, C. Y., S. El-Sharnouby, A. Chatzipli, S. Russell, S. W. Choo et al., 2016 Roles of cofactors and chromatin accessibility in Hox protein target specificity. Epigenetics Chromatin 9: 1.

Berloco, M., L. Fanti, A. Breiling, V. Orlando, and S. Pimpinelli, 2001 The maternal effect gene, abnormal oocyte (abo), of Drosophila melanogaster encodes a specific negative regulator of histones. Proc Natl Acad Sci U S A 98: 12126–12131.

Bongartz, P., and S. Schloissnig, 2018 Deep repeat resolution—the assembly of the Drosophila Histone Complex. Nucleic Acids Research 47: e18–e18.

Bulchand, S., S. D. Menon, S. E. George, and W. Chia, 2010 Muscle wasted: a novel component of the Drosophila histone locus body required for muscle integrity. Journal of Cell Science 123: 2697–2707.

Cai, W., C. Wang, Y. Li, C. Yao, L. Shen et al., 2014 Genome-wide analysis of regulation of gene expression and H3K9me2 distribution by JIL-1 kinase mediated histone H3S10 phosphorylation in Drosophila. Nucleic Acids Res 42: 5456–5467.

Chari, S., H. Wilky, J. Govindan, and A. A. Amodeo, 2019 Histone concentration regulates the cell cycle and transcription in early development. Development 146: dev177402.

Crayton, M. E., C. E. Ladd, M. Sommer, G. Hampikian, and L. D. Strausbaugh, 2004 An organizational model of transcription factor binding sites for a histone promoter in D. melanogaster. In Silico Biol 4: 537–548.

Daneshvar, K., A. Khan, and J. M. Goodliffe, 2011 Myc Localizes to Histone Locus Bodies during Replication in Drosophila. PLOS ONE 6: e23928.

Docquier, F., O. Saget, F. Forquignon, N. B. Randsholt, and P. Santamaria, 1996 The multi sex combs gene of Drosophila melanogaster is required for proliferation of the germline. Rouxs Arch Dev Biol 205: 203–214.

Doiguchi, M., T. Nakagawa, Y. Imamura, M. Yoneda, M. Higashi et al., 2016 SMARCAD1 is an ATP-dependent stimulator of nucleosomal H2A acetylation via CBP, resulting in transcriptional regulation. Sci Rep 6: 20179.

Duronio, R. J., and W. F. Marzluff, 2017 Coordinating cell cycle-regulated histone gene expression through assembly and function of the Histone Locus Body. RNA Biol 14: 726–738.

Feng, S., C. Rastogi, R. Loker, W. J. Glassford, H. Tomas Rube et al., 2022 Transcription factor paralogs orchestrate alternative gene regulatory networks by context-dependent cooperation with multiple cofactors. Nat Commun 13: 3808.

Follmer, N. E., A. H. Wani, and N. J. Francis, 2012 A polycomb group protein is retained at specific sites on chromatin in mitosis. PLoS Genet 8: e1003135.

Gramates, L. S., J. Agapite, H. Attrill, B. R. Calvi, M. A. Crosby et al., 2022 FlyBase: a guided tour of highlighted features. Genetics 220: iyac035.

Herz, H.-M., M. Mohan, A. S. Garrett, C. Miller, D. Casto et al., 2012 Polycomb repressive complex 2-dependent and -independent functions of Jarid2 in transcriptional regulation in Drosophila. Mol Cell Biol 32: 1683–1693.

Isogai, Y., S. Keles, M. Prestel, A. Hochheimer, and R. Tjian, 2007 Transcription of histone gene cluster by differential core-promoter factors. Genes Dev. 21: 2936–2949.

Jimeno-González, S., L. Payán-Bravo, A. M. Muñoz-Cabello, M. Guijo, G. Gutierrez et al., 2015 Defective histone supply causes changes in RNA polymerase II elongation rate and cotranscriptional pre-mRNA splicing. Proceedings of the National Academy of Sciences 112:14840–14845.

Kang, H., K. A. McElroy, Y. L. Jung, A. A. Alekseyenko, B. M. Zee et al., 2015 Sex comb on midleg (Scm) is a functional link between PcG-repressive complexes in Drosophila. Genes Dev. 29: 1136–1150.

Kaya-Okur, H. S., S. J. Wu, C. A. Codomo, E. S. Pledger, T. D. Bryson et al., 2019 CUT&Tag for efficient epigenomic profiling of small samples and single cells. Nat Commun 10: 1930.

Kaye, E. G., M. Booker, J. V. Kurland, A. E. Conicella, N. L. Fawzi et al., 2018 Differential Occupancy of Two GA-Binding Proteins Promotes Targeting of the Drosophila Dosage Compensation Complex to the Male X Chromosome. Cell Rep 22: 3227–3239.

Kellner, W. A., K. Van Bortle, L. Li, E. Ramos, N. Takenaka et al., 2013 Distinct isoforms of the Drosophila Brd4 homologue are present at enhancers, promoters and insulator sites. Nucleic Acids Res 41: 9274–9283.

Kemp, J. P., X.-C. Yang, Z. Dominski, W. F. Marzluff, and R. J. Duronio, 2021 Superresolution light microscopy of the Drosophila histone locus body reveals a core–shell organization associated with expression of replication–dependent histone genes. MBoC 32: 942–955.

Koenecke, N., J. Johnston, B. Gaertner, M. Natarajan, and J. Zeitlinger, 2016 Genome-wide identification of Drosophila dorso-ventral enhancers by differential histone acetylation analysis. Genome Biology 17: 196.

Koreski, K. P., L. E. Rieder, L. M. McLain, W. F. Marzluff, and R. J. Duronio, 2020 Drosophila Histone Locus Body assembly and function involves multiple interactions. bioRxiv 2020.03.16.994483.

Koromila, T., F. Gao, Y. Iwasaki, P. He, L. Pachter et al., 2020 Odd-paired is a pioneer-like factor that coordinates with Zelda to control gene expression in embryos (K. Struhl, O. Hobert, & E. Clark, Eds.). eLife 9: e59610.

Kribelbauer, J. F., R. E. Loker, S. Feng, C. Rastogi, N. Abe et al., 2020 Context-Dependent Gene Regulation by Homeodomain Transcription Factor Complexes Revealed by Shape-Readout Deficient Proteins. Molecular Cell 78: 152–167.e11.

Kyrchanova, O., N. Klimenko, N. Postika, A. Bonchuk, N. Zolotarev et al., 2021 Drosophila architectural protein CTCF is not essential for fly survival and is able to function independently of CP190. Biochimica et Biophysica Acta (BBA) - Gene Regulatory Mechanisms 1864: 194733.

Langmead, B., and S. L. Salzberg, 2012 Fast gapped-read alignment with Bowtie 2. Nat Methods 9: 357–359.

Larschan, E., A. A. Alekseyenko, W. R. Lai, P. J. Park, and M. I. Kuroda, 2006 MSL complex associates with clusters of actively transcribed genes along the Drosophila male X chromosome. Cold Spring Harb Symp Quant Biol 71: 385–94.

Larschan, E., M. M. Soruco, O. K. Lee, S. Peng, E. Bishop et al., 2012 Identification of chromatin-associated regulators of MSL complex targeting in Drosophila dosage compensation. PLoS Genet 8: e1002830.

Li, Z., M. R. Johnson, Z. Ke, L. Chen, and M. A. Welte, 2014 Drosophila lipid droplets buffer the H2Av supply to protect early embryonic development. Curr Biol 24: 1485–1491.

Li, Z., K. Thiel, P. J. Thul, M. Beller, R. P. Kühnlein et al., 2012 Lipid droplets control the maternal histone supply of Drosophila embryos. Curr Biol 22: 2104–2113.

Liu, J.-L., C. Murphy, M. Buszczak, S. Clatterbuck, R. Goodman et al., 2006 The Drosophila melanogaster Cajal body. J Cell Biol 172: 875–884.

Marzluff, W. F., P. Gongidi, K. R. Woods, J. Jin, and L. J. Maltais, 2002 The human and mouse replication-dependent histone genes. Genomics 80: 487–98.

Matera, A. G., M. Izaguire-Sierra, K. Praveen, and T. K. Rajendra, 2009 Nuclear Bodies: Random Aggregates of Sticky Proteins or Crucibles of Macromolecular Assembly? Developmental Cell 17: 639–647.

McKay, D. J., S. Klusza, T. J. Penke, M. P. Meers, K. P. Curry et al., 2015 Interrogating the function of metazoan histones using engineered gene clusters. Dev Cell 32: 373–86.

Ozawa, N., H. Furuhashi, K. Masuko, E. Numao, T. Makino et al., 2016 Organ identity specification factor WGE localizes to the histone locus body and regulates histone expression to ensure genomic stability in Drosophila. Genes to Cells 21: 442–456.

Ramalingam, V., M. Natarajan, J. Johnston, and J. Zeitlinger, 2021 TATA and paused promoters active in differentiated tissues have distinct expression characteristics. Molecular Systems Biology 17: e9866.

Ravindranath, A., and K. M. Cadigan, 2014 Structure-Function Analysis of the C-clamp of TCF/Pangolin in Wnt/ß-catenin Signaling. PLOS ONE 9: e86180.

Regnard, C., T. Straub, A. Mitterweger, I. K. Dahlsveen, V. Fabian et al., 2011 Global Analysis of the Relationship between JIL-1 Kinase and Transcription. PLOS Genetics 7: e1001327.

Rezsohazy, R., A. J. Saurin, C. Maurel-Zaffran, and Y. Graba, 2015 Cellular and molecular insights into Hox protein action. Development 142: 1212–1227.

Rhee, H. S., and B. F. Pugh, 2012 ChIP-exo Method for Identifying Genomic Location of DNA-Binding Proteins with Near-Single-Nucleotide Accuracy. Current Protocols in Molecular Biology 100: 21.24.1–21.24.14.

Rieder, L. E., K. P. Koreski, K. A. Boltz, G. Kuzu, J. A. Urban et al., 2017 Histone locus regulation by the Drosophila dosage compensation adaptor protein CLAMP. Genes Dev 31: 1494–1508.

Robinson, J. T., H. Thorvaldsdóttir, W. Winckler, M. Guttman, E. S. Lander et al., 2011 Integrative Genomics Viewer. Nat Biotechnol 29: 24–26.

Saget, O., F. Forquignon, P. Santamaria, and N. B. Randsholt, 1998 Needs and targets for the multi sex combs gene product in Drosophila melanogaster. Genetics 149: 1823–1838.

Salzler, H. R., D. C. Tatomer, P. Y. Malek, S. L. McDaniel, A. N. Orlando et al., 2013 A sequence in the Drosophila H3-H4 Promoter triggers histone locus body assembly and biosynthesis of replication-coupled histone mRNAs. Dev Cell 24: 623–34.

Schmidt, C. A., L. J. Hodkinson, H. S. Comstra, and L. E. Rieder, 2022 A cost-free CURE: Using bioinformatics to identify DNA-binding factors at a specific genomic locus. 2022.10.21.513244.

Shevtsov, S. P., and M. Dundr, 2011 Nucleation of nuclear bodies by RNA. Nat Cell Biol 13: 167–173.

Shlyueva, D., A. C. A. Meireles-Filho, M. Pagani, and A. Stark, 2016 Genome-Wide Ultrabithorax Binding Analysis Reveals Highly Targeted Genomic Loci at Developmental Regulators and a Potential Connection to Polycomb-Mediated Regulation. PLoS One 11: e0161997.

Stephenson, R. A., J. M. Thomalla, L. Chen, P. Kolkhof, R. P. White et al., 2021 Sequestration to lipid droplets promotes histone availability by preventing turnover of excess histones. Development 148: dev199381.

Straub, T., A. Zabel, G. D. Gilfillan, C. Feller, and P. B. Becker, 2013 Different chromatin interfaces of the *Drosophila* dosage compensation complex revealed by high-shear ChIP-seq. Genome Res. 23: 473–485.

Szklarczyk, D., A. L. Gable, D. Lyon, A. Junge, S. Wyder et al., 2019 STRING v11: protein-protein association networks with increased coverage, supporting functional discovery in genome-wide experimental datasets. Nucleic Acids Res 47: D607–D613.

Tatomer, D. C., E. Terzo, K. P. Curry, H. Salzler, I. Sabath et al., 2016 Concentrating pre-mRNA processing factors in the histone locus body facilitates efficient histone mRNA biogenesis. Journal of Cell Biology 213: 557–570.

Terzo, E. A., S. M. Lyons, J. S. Poulton, B. R. S. Temple, W. F. Marzluff et al., 2015 Distinct self-interaction domains promote Multi Sex Combs accumulation in and formation of the Drosophila histone locus body. Mol Biol Cell 26: 1559–1574.

The Galaxy Community, 2022 The Galaxy platform for accessible, reproducible and collaborative biomedical analyses: 2022 update. Nucleic Acids Research 50: W345–W351.

THE MODENCODE CONSORTIUM, S. Roy, J. Ernst, P. V. Kharchenko, P. Kheradpour et al., 2010 Identification of Functional Elements and Regulatory Circuits by Drosophila modENCODE. Science 330: 1787–1797.

Vivekanand, P., 2018 Lessons from Drosophila Pointed, an ETS family transcription factor and key nuclear effector of the RTK signaling pathway. Genesis 56: e23257.

Wang, C. I., A. A. Alekseyenko, G. LeRoy, A. E. H. Elia, A. A. Gorchakov et al., 2013 Chromatin proteins captured by ChIP-mass spectrometry are linked to dosage compensation in Drosophila. Nat Struct Mol Biol 20: 202–209.

Webber, J. L., J. Zhang, A. Massey, N. Sanchez-Luege, and I. Rebay, 2018 Collaborative repressive action of the antagonistic ETS transcription factors Pointed and Yan fine-tunes gene expression to confer robustness in Drosophila. Development 145: dev165985.

White, A. E., B. D. Burch, X. C. Yang, P. Y. Gasdaska, Z. Dominski et al., 2011 Drosophila histone locus bodies form by hierarchical recruitment of components. J Cell Biol 193: 677–94.

Wolle, D., F. Cleard, T. Aoki, G. Deshpande, P. Schedl et al., 2015 Functional Requirements for Fab-7 Boundary Activity in the Bithorax Complex. Molecular and Cellular Biology 35: 3739–3752.

Xie, M., S. Comstra, C. Schmidt, L. Hodkinson, and L. E. Rieder, 2022a Max is likely not at the Drosophila histone locus. 2022.09.11.507040.

Xie, M., L. J. Hodkinson, H. S. Comstra, P. P. Diaz-Saldana, H. E. Gilbonio et al., 2022b MSL2 targets histone genes in Drosophila virilis. 2022.12.14.520423.

Yang, X., I. Sabath, L. Kunduru, A. J. van Wijnen, W. F. Marzluff et al., 2014 A conserved interaction that is essential for the biogenesis of histone locus bodies. J Biol Chem 289: 33767–33782.

Yusufova, N., A. Kloetgen, M. Teater, A. Osunsade, J. M. Camarillo et al., 2021 Histone H1 loss drives lymphoma by disrupting 3D chromatin architecture. Nature 589: 299–305.

