## Supplemental Table 1 for "A bioinformatics screen reveals Hox and chromatin remodeling factors at the *Drosophila* histone locus"

| Candidate | Category | Rationale | Tissue/Timing | Localization | Region |
|---|---|---|---|---|---|
| **Abd-A**<br>Abdominal A | Early development transcription factor | Continuous expression post 0-2hour developmental stage, with highest RNA expression levels between 2-4hrs (THE MODENCODE CONSORTIUM *et al.* 2010; Gramates *et al.* 2022) | Kc cells | Yes | *H3/H4* promoter |
| **Abd-B**<br>abdominal B | Early development transcription factor | Continuous expression post 0-2hour developmental stage, with highest RNA expression levels between 2-4hrs (THE MODENCODE CONSORTIUM *et al.* 2010; Gramates *et al.* 2022) | Kc cells | Yes | *H3/H4* promoter |
| **Antp**<br>Antennapedia | Early development transcription factor | Expressed in the early embryo (THE MODENCODE CONSORTIUM *et al.* 2010; Gramates *et al.* 2022) | Imaginal wing disk | | |
| **Ndf (CG4747)**<br>Nucleosome-destabilizing factor | Dosage compensation associated factor | H3K36me3-binding protein that is important for MSLc localization (Wang *et al.* 2013) | larval | | |
| **CP190**<br>Centrosomal protein 190kD | Chromatin structure/remodeler | Insulates centromieric heterochromatin (Bag *et al.* 2019) | Kc/S2 cells | | |
| **Exd**<br>extradenticle | Early development transcription factor | Expressed in the early embryo (THE MODENCODE CONSORTIUM *et al.* 2010; Gramates *et al.* 2022) | 5-6 days wing | | |
| **fs(1)h**<br>female sterile (1) homeotic | Chromatin structure/remodeler | short isoform binds at promoters and enhancers and long isoform binds at chromatin insulators<br>(Kellner *et al.* 2013) | Kc cells | Yes (long isoform only) | *H3/H4* promoter<br>*H2A/H2B* promoter |
| **Gcn5** | Chromatin structure/remodeler | Acetyltransferase, critical for oogenesis and morphogenesis (Ali *et al.* 2017) and associates with insulators (Bag *et al.* 2019) | Kc cells | | |
| **Hr78**<br>Hormone-receptor-like in 78 | Early development transcription factor | Continuous expression throughout embryogenesis and during metamorphosis (THE MODENCODE CONSORTIUM *et al.* 2010; Gramates *et al.* 2022) | 8-16h embryos | Yes | *H3/H4* promoter |
| **Hth**<br>homothorax | Early development transcription factor | Expressed in the early embryo (THE MODENCODE CONSORTIUM *et al.* 2010; Gramates *et al.* 2022) | Imaginal wing disc | | |
| **JIL-1** | Dosage compensation/ X-chromosome associated factor | Kinase phosphorylation of H3S10<br>Enriched on X chromosome for dosage compensation<br>Serine 10 mark prevents heterochromatin spreading<br>(Regnard *et al.* 2011; Cai *et al.* 2014; Albig *et al.* 2019) | 3$^{rd}$ instar larvae | Yes | *H2A/H2B* promoter |
| **M1BP**<br>Motif 1 Binding Protein | Chromatin structure/remodeler | Interacts with TRF2 and related to insulator activity by associating with CP190 (Bag *et al.* 2021*)* | Kc/S2 cells | | |
| **MSL1**<br>Male specific Lethal 1 | Dosage compensation factor | Associates with the known HLB factor CLAMP (Larschan *et al.* 2012; Kaye *et al.* 2018) | S2 cells | | |
| **Nej**<br>Nejire | Hit from previous screen for HLB factors<br>(White *et al.* 2011) | Acetyltransferase and early developmental transcription factor | 2-4 hr embryos<br>S2 cells | Yes (embryos only) | *H3/H4* promoter<br>*H2A/H2B* promoter |
| **Opa**<br>Odd Paired | Early development transcription factor | Continuous expression post 0-2hour developmental stage, with highest RNA expression levels between 2-4hr when ZGA/ histone gene expression is high (THE MODENCODE CONSORTIUM *et al.* 2010; Gramates *et al.* 2022) | 3h embryo<br>4h embryo | | |
| **Pan**<br>Pangolin | Early development transcription factor | Expressed in the early embryo/early in development (THE MODENCODE CONSORTIUM *et al.* 2010; Gramates *et al.* 2022) | 0-8h embryo | | |
| **Pnt**<br>Pointed | Hit from previous screen for HLB factors<br>(White *et al.* 2011) | Early development transcription factor | Stage 11 embryo | | |

| | | | | | |
|---|---|---|---|---|---|
| **Psc**<br>Posterior sex combs | Chromatin structure/remodeler | Polycomb member (Follmer *et al.* 2012) | S2 cells | | |
| **Scm**<br>Sex comb on midleg | HLB-associated factor | Genetically interacts with known HLB factor Mxc (Docquier *et al.* 1996; Saget *et al.* 1998) | Embryo 12-24h<br>S2 cells | | |
| **su(z)12**<br>suppressor of zeste 12 | Chromatin structure/remodeler | Polycomb repressive complex member (Herz *et al.* 2012)<br>Highest RNA expression levels during 0-2h and 4-8h development stages (THE MODENCODE CONSORTIUM *et al.* 2010; Gramates *et al.* 2022) | 3rd instar larvae | | |
| **TAF1**<br>TBP-associated factor 1 | General transcription factor | TATA-box-binding protein known to associate with TBP (Baumann and Gilmour 2017) | S2 cells | Yes | *H3/H4* promoter |
| **TFIIB**<br>Transcription Factor II B | General transcription factor | TATA-box binding protein complex member, known to associate with TBP (Ramalingam *et al.* 2021) | OregonR Embyos | Yes | *H3/H4* promoter<br>*H2A/H2B* promoter |
| **TFIIF**<br>Transcription Factor II F | General transcription factor | TATA-box binding protein complex member, known to associate with TBP (Ramalingam *et al.* 2021) | OregonR Embyos | Yes | *H3/H4* promoter<br>*H2A/H2B* promoter<br>*H1* promoter |
| **TRF2**<br>TATA box binding protein-related factor 2 | General transcription factor | TATA-less promoter binding activity at *H1* promoter (Isogai *et al.* 2007) | S2 cells | Yes | *H1* promoter |
| **Ubx**<br>Ultrabithorax | Early development transcription factor | Continuous expression post 0-2hour developmental stage, with highest RNA expression levels between 2-4hrs (THE MODENCODE CONSORTIUM *et al.* 2010; Gramates *et al.* 2022) | Kc cells,<br>imaginal wing disc embryos | Yes (all) | *H3/H4* promoter |
